# Environmental filtering in marine fish communities is weakened during severe hypoxia season

**DOI:** 10.1101/2021.04.29.442042

**Authors:** Naoto Shinohara, Yuki Hongo, Momoko Ichinokawa, Shota Nishijima, Shuhei Sawayama, Hiroaki Kurogi, Yasuyuki Uto, Hisanori Mita, Mitsuhiro Ishii, Akane Kusano, Seiji Akimoto

## Abstract

Compositional variation among local communities reflects the differences in abiotic environments, though the extent of the influence varies in natural systems. Generally, the environmental filtering is expected to be strong if the environmental gradient encompasses severe habitats where species sorting is highly likely to work. However, this hypothesis has rarely been tested in dynamic systems, where the higher dispersal ability of species allows them to easily respond to stressful environments. Here, with the dynamics of fish communities in a Japanese bay revealed by environmental DNA analyses as a model case, we examined how harmful seasonal hypoxia (low concentration of oxygen in bottom waters in summer) affected the strength of environmental filtering. We found that, in summer, dissolved oxygen (DO) concentration was significantly low and fish species richness decreased in the bottom water compared to the surface, suggesting that the organisms were adversely affected by the hypoxia. At the same time, the between-depths heterogeneity in DO concentration was larger, but species composition was less divergent and the influence of DO on species composition appeared weaker during summer. These results imply that environmental filtering is weakened when the bottom water was characterized by extremely severe environments. Furthermore, there was a shift in the species occurrence from bottom to surface waters in summer that was consistent across species, suggesting that the extremely severe hypoxia adversely affected fish species irrespective of their identity. These results collectively suggest that environmental filtering is weaker during summer despite more severity and heterogeneity in environments, most likely because individual movements to avoid unpreferable environments in the bottom waters occurred quasi-neutrally. By providing evidence against the prevailing understanding that environmental filtering strongly works in severe environments, these findings invoke further investigation on how the filtering acts in various conditions.

## Introduction

Understanding the processes structuring local communities has been a central challenge in ecology (Hutchinson 1959, HilleRisLambers et al. 2012). One of the primary determinants of local communities is the abiotic environment; species composition is thought to deterministically align with environmental gradients as determined by each species’ requirements for some habitats or abiotic conditions (Grinnell 1917, Tilman 1982), a concept commonly referred to as ‘environmental filtering’. However, it has also been acknowledged that more stochastic processes such as random movements of individuals among localities influence the community structure (Sale 1978, Hubbell 2001). Therefore, species composition can vary even if the localities are similar in their environmental conditions. Acknowledging that both deterministic and stochastic processes act in concert in nature, much research on community assembly has been devoted to understanding to what extent and in what situation the deterministic (or stochastic) community assembly matters (Cottenie 2005, Chase and Myers 2011).

Growing body of literature suggests that deterministic assembly plays an important role in more abiotically severe habitats (Chase 2007, Lepori and Malmqvist 2009, Guo et al. 2014), because the stressful environment filters the potential colonizers. The reduced potential species pool would hamper the importance of stochastic assembly by reducing the among-habitats variation in species composition and the probability of alternative states (Chase 2007). Therefore, environmental filtering is expected to be more influential when the environmental gradient encompasses severe conditions that strongly sort species. However, there exist a couple of studies providing opposing results that stochasticity is amplified in stressful environments (Lepori and Malmqvist 2009, Kim 2019). For example, although the deterministic control is weak in streams without disturbance, it is also so in highly disturbed habitats because the disturbance can induce extinction of individuals irrespective of species traits (Lepori and Malmqvist 2009). Therefore, whether environmental severity intensifies or dampens the deterministic control of environments is still an open question in community ecology. Furthermore, as previous studies have focused on organisms with limited or passive dispersal abilities (Chase 2007, Guo et al. 2014), knowledge is particularly scarce in dynamic communities characterized by higher dispersal ability of species. If species can perceive and easily respond to environmental severity, the filtering may act more precisely. Otherwise, it is also probable that species actively avoid unpreferable, though tolerable, environments, which dampens the species sorting by environments.

In marine ecosystems, one of the environmental variables that most drastically influence community structures is hypoxia or the decline in dissolved oxygen (DO; Diaz and Rosenberg 1995, 2008, Breitburg et al. 2018). One of the typical forms of hypoxia is seasonal: In summer, water warming forms the stratification of water columns, which reduce the DO concentration in the bottom water to a critical level. As oxygen is essential to organisms, the decreased DO adversely affects demersal organisms’ performance (Levin 2003, Hrycik et al. 2017). Therefore, the bottom hypoxia is expected to serve as a strong environmental filter by sorting out the species that are vulnerable to the lower DO conditions (Levin 2003, Kodama and Horiguchi 2011), which aligns with the idea that severe abiotic conditions intensify the deterministic assembly (Chase 2007, Lepori and Malmqvist 2009, Guo et al. 2014). On the contrary, if the bottom hypoxia was so intense that all species suffer from or avoid it to a similar extent, species composition would no longer be correlated with the surface-bottom gradient in DO, making environmental filtering weak.

The objective of this study is to determine how extremely severe conditions alter the strength of environmental filtering in a dynamic system. We focused on the influence of bottom hypoxia in summer on fish communities in Tokyo Bay, one of the largest bay areas in Japan, as a model case. In this study system, it is already known that the water stratification in summer causes hypoxia (Yasui et al. 2016) which severely affects organisms in the bottom waters (Kodama and Horiguchi 2011). We have two alternative hypotheses: (1) filtering by DO is stronger when the bottom hypoxia is more intense (in summer) as fish species are different in their ability to tolerate lower oxygen concentration. This predicts that explanatory power of DO for fish species composition is high in summer because the species composition is clearly separated between oxygen-limited bottom and oxygen-rich surface. Alternatively, (2) the filtering is weak in summer because most of the fish species similarly suffer from or avoid the lower DO in the bottom. This alternative hypothesis predicts that the explanatory power of DO is weak and surface-bottom compositional differentiation is less evident in summer because the frequency of occurrence in the bottom water is decreased irrespective of species identity. To this end, we took advantage of the availability of data of seasonal dynamics of fish communities in Tokyo Bay revealed by environmental DNA (eDNA) analyses (Hongo et al. 2021) which has emerged as a powerful and efficient tool for monitoring biodiversity especially in aquatic ecosystems (Bohmann et al. 2014; Deiner et al. 2017). By extracting genetic materials shed from organisms from environmental samples (e.g., water and soils), this technique allows for the non-invasive assessment of biodiversity, with a detection ability that is comparable to that of traditional sampling methods (Thomsen et al. 2012; Sigsgaard et al. 2017; Yamamoto et al. 2017). Importantly, owing to the short persistence (from a few hours [Murakami et al. 2019] to several days [Thomsen et al. 2012]) and low diffusion rate (less than 100 m; Port et al. 2016; Murakami et al. 2019) of eDNA in seawater, this technique enables the estimation of contemporary and local species composition (Yamamoto et al. 2016). Taking advantage of this technique allowed us to sample fish in both bottom and surface waters, at a high frequency throughout a single year.

## Materials and methods

### Water sampling and eDNA sequencing

Water sampling was conducted monthly at 14 sites across Tokyo Bay, Japan (approximately 1380 km^2^, 35°30’ N, 139°50’ E, Fig. S1) from January to December 2019. At each sampling location, 1 L of water was collected from the surface and bottom waters. Bottom depth varied among sites from approximately 6 to 70 m. Water samples were filtered and frozen onboard a ship. The frozen filters were then transferred to and stored in the laboratory. The extraction of eDNA and the construction of amplicon sequencing variants (ASVs) were performed as described by Hongo et al. (2021) using MiFish universal primers (Miya et al. 2015). The constructed ASVs were assigned to fish species using Blastn against the MitoFish database (Sato et al. 2018). As the correlation between eDNA concentration and abundance of target fish species is variable under natural conditions (Yates et al. 2019), the count data of ASVs was replaced to presence/absence data. The eDNA sequence data are available at the DNA Data Bank of Japan Sequence Read Archive under the accession number DRA010909.

Concentration of DO was measured at the same time as the water sampling. As expected, we found that DO of the bottom water was lower and the value was more divergent between surface and bottom during summer (Fig. 1). Specifically, from June to September, the minimum value of DO concentration among samples was lower than 2 mg/L, a common threshold of hypoxia in previous studies (Pihl et al. 1991, Hrycik et al. 2017, Breitburg et al. 2018). Following the established threshold in the literature, we determined these four months as “hypoxia season” and other months as “normoxia” (normal levels of oxygen). We note that even in the hypoxia season, the DO values considerably varied among samples, with many of them being higher than the threshold (Fig. 1). Nevertheless, we classified all samples of the months when the minimum value falls below the threshold as the hypoxia season. This is because our hypothesis was that the explanatory power of DO for fish species composition is strong (or weak) when the environmental gradient encompasses the severe conditions, and we therefore needed to analyze the samples obtained in the same environmental gradient as one unit.

**Figure 1.**
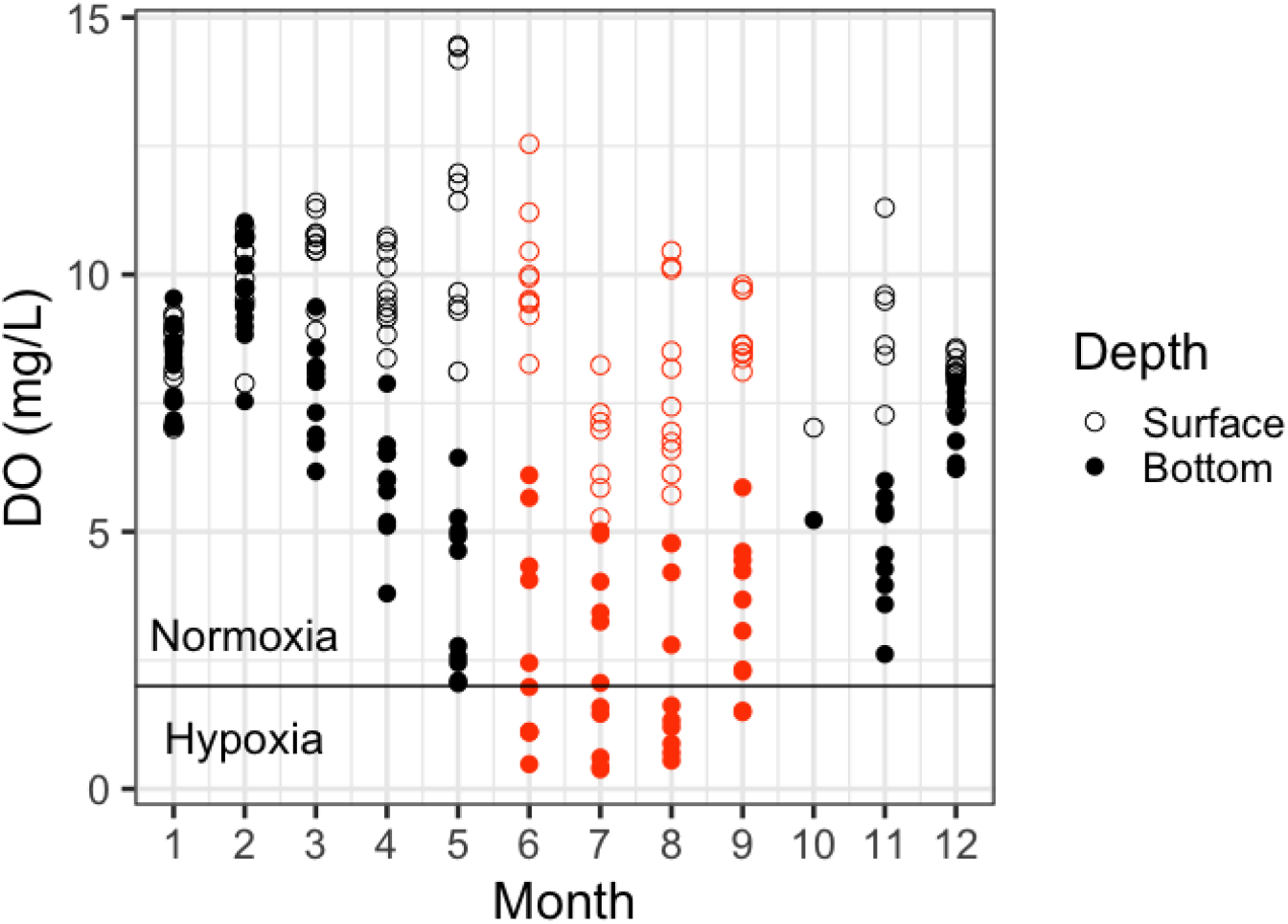
Temporal trend of concentration of dissolved oxygen (DO) of each sample. Open and filled circles represent samples in surface and bottom waters, respectively. The minimum value of DO concentration was lower than the common threshold (2 mg/L) from June to September, the period we determined as hypoxia season (coloured with orange).

### Data analysis

To keep the dataset free from unexpected biases such as sequencing errors or contamination from rivers or nearby fish markets, we filtered out the species that did not meet the following two criteria from further analyses: (1) those whose names were not matched during the search in Fishbase, a global database of fish (Froese and Pauly 2019; https://www.fishbase.se/), and (2) those whose habitats were known to be out of Tokyo Bay based on information from a range of studies (Nakabo 2002, Kouno et al. 2011). The first criterion mainly filtered out hybrid species, while the second one, which was based on each fish species’ distribution in Japan and observation records in Tokyo Bay, removed fish species known to be freshwater or endemic in other regions.

All statistical analyses were performed using R version 4.0.2 (R Core Team 2020) and the figures were illustrated using the *ggplot2* package (Wickham 2016). First, to examine the overall effects of lower DO on fish communities, we constructed generalized linear models (GLMs) that modelled the fish species richness as a function of DO with Poisson distribution (log link). The models were built for the bottom and surface samples separately. Second, the explanatory power of DO for the fish species composition was evaluated using distance-based redundancy analyses (db-RDA, Legendre and Anderson 1999) that takes the Jaccard dissimilarity matrix (based on presence-absence data) as a response and DO value as an explanatory variable. The analyses were conducted separately for each month, and the explanatory power was evaluated as the R^2^ value that was adjusted for the number of variables (Peres-Neto et al. 2006) using the *RsquareAdj* function in the *vegan* package (Oksanen et al. 2019). The R^2^ values were compared between hypoxia and normoxia months by Wilcoxon’s rank sum exact test. Third, to look at the surface-bottom divergence of species composition, we conducted nonmetric multidimensional scaling (NMDS) on the whole species composition data (n=281) using the *metaMDS* function. This allows the compositional dissimilarity to be interpreted in a low-dimensional space by positioning similar data nearby. For the NMDS analysis, we used the Jaccard dissimilarity index on the presence-absence dataset and three dimensions (k=3) to facilitate convergence. For the interpretation, we plotted values of the first two MDS axes in the figure and illustrated ellipses of 80% confidence level for multivariate normal distributions for each season (hypoxia or normoxia) and depth (surface or bottom). Difference in the species composition among seasons and depths was tested using permutational multivariate analysis of variance (PERMANOVA; Anderson 2001, the *adonis* function). In addition, we included the interaction term between season and depth in the factorial design of the PERMANOVA to test the difference in the degree of surface-bottom divergence of species composition between seasons. Since the number of sampling sites was small (n=2) due to logistical problems, we did not conduct the variation partitioning analysis and PERMANOVA for the October data.

Finally, to examine whether the responses to bottom hypoxia are species-specific, we compared the frequency of occurrence of fish species among different seasons (hypoxia or normoxia) and depths (surface or bottom), using generalized linear mixed models (GLMMs) with species-specific random intercept and coefficients. The models were implemented using the *glmmTMB* function in the *glmmTMB* package (Brooks et al. 2017). The response variable was the presence probability of species *i* in a sample *s* (*p_i,s_*), and the explanatory variables were seasons, depths, and their interaction:

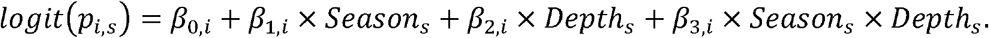

Note that the season and depth were transformed into dummy variables here (hypoxia: 0, normoxia: 1, and surface: 0, bottom: 1). To assure sufficient statistical power, we restricted this analysis to the frequently observed species, that is, top 10% most frequently observed species (17 out of 168 species). Therefore, the dataset used in this GLMM consisted of 3842 presence or absence records (17 species x 226 samples). The species-specific intercept (*β*_0,*i*_) and coefficients (*β*_1,*i*_, *β*_2,*i*_, *β*_3,*i*_) were modelled to follow normal distributions, reflecting that the baseline of occurrence probability, the effects of seasons, depth and their interaction on the probability may differ among species. Significance of the estimated fixed coefficients suggests that the effects of the explanatory variables were consistent while accounting for the difference among species. In light of our hypothesis, the significantly positive coefficient of the fixed effect of the interaction term implies that the relative occurrence in the bottom over surface water in normoxia was higher on average across species, suggesting that species generally suffered from bottom hypoxia. Conversely, if the coefficient of the interaction was not significant, the response to bottom hypoxia is not evident at the average level among species. To further test the hypothesis on species-specific responses to bottom hypoxia, we additionally constructed a GLMM that was similar to the previous one except for that the coefficient of the interaction term (*β*_3,*i*_) was modelled as a fixed effect (*β*_3_). We compared the fittings of the two GLMMs by the likelihood ratio test using the *anova* function, which informs us whether modelling the species-specific responses to bottom hypoxia improves the model. If the test indicated no significant differences between them, we conclude that the effect of season-depth interaction can be modelled as common across species.

## Results

We obtained 308 water samples for the eDNA extraction, and gene amplification were successful for 281 samples. Among these, corresponding DO data were available for 225 samples (n = 22, 22, 22, 21, 20, 20, 18, 21, 20, 2, 15, and 22 from January to December, respectively). A total of 220 fish species were detected in the 225 samples. Of these, our first criterion filtered out 12 species, and the second criterion eliminated 40 species, resulting in 168 species retained in the subsequent analyses (Table S1).

The average species richness per sample was higher in the bottom layer (mean: 9.73, standard deviation: 4.78) than in the surface layer (mean: 6.89, standard deviation: 3.20) (Fig. S2). The GLMs that modelled fish species richness as functions of DO indicate that the species richness in the bottom water was positively correlated with DO (GLM: coefficient = 0.037; *p* < 0.001, Fig. 2), while that in the surface water decreased with DO (coefficient = −0.085; *p* < 0.001).

**Figure 2.**
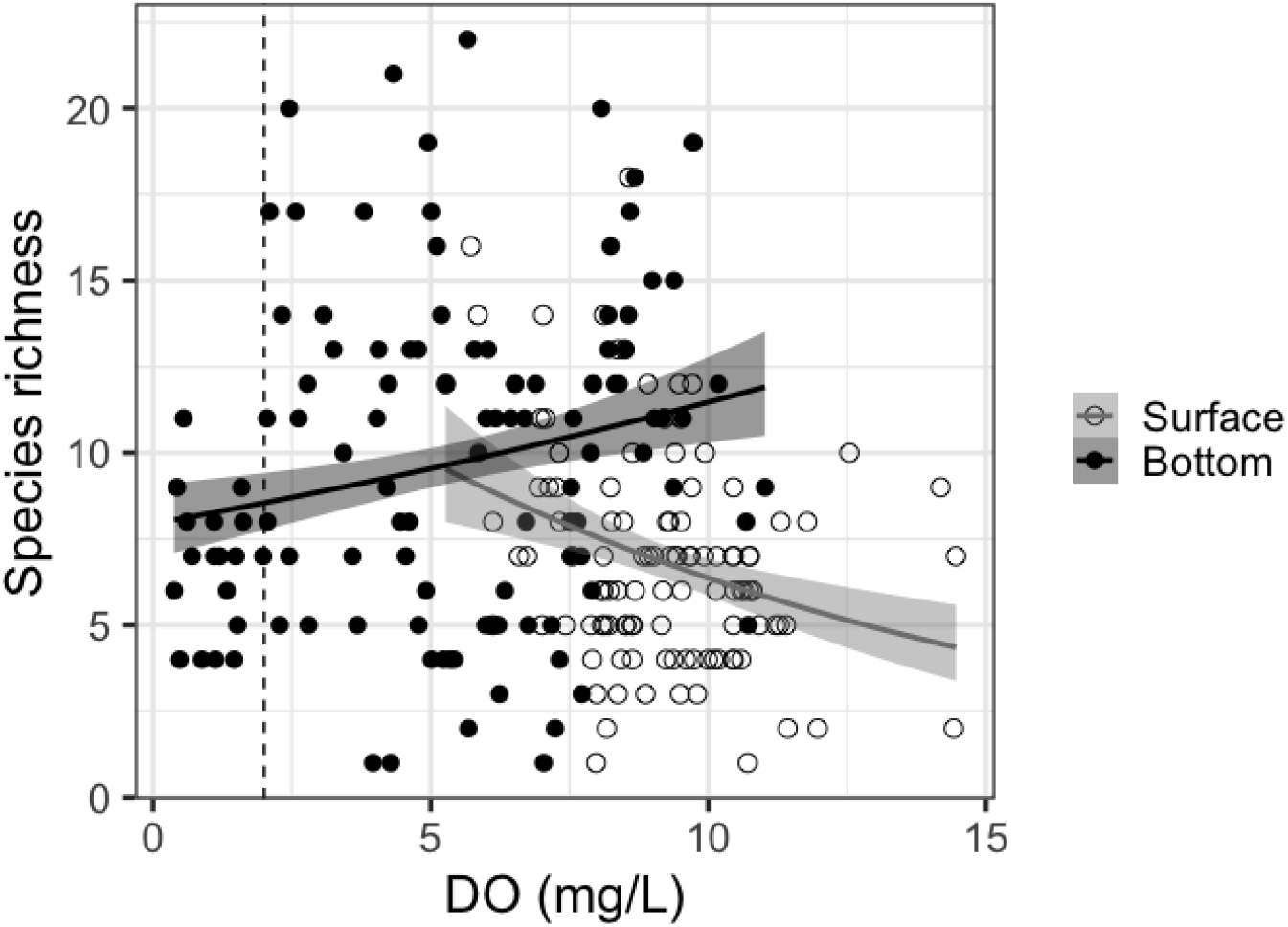
Relationship between dissolved oxygen (DO) concentration and fish species richness. Open and filled circles represent samples in surface and bottom waters, respectively. The vertical broken line indicates the threshold for the hypoxia (2 mg/L). The lines were drawn based on the generalized linear models separately for the depths, after confirming the statistical significance (*p* < 0.05) of the slopes (see the main text).

The variation partitioning analyses suggested that DO value explained 6.4% of the variation of fish species composition on average across months (n = 11). Note that the analysis was not conducted here for the October data because of small sample size. The explanatory power was 4.2% on average in hypoxia (n = 4), whereas that was 7.7% in normoxia (n = 7). Wilcoxon’s rank sum exact test indicated that the explanatory power was significantly higher in normoxia (W = 3; *p* = 0.040, Fig. S3).

The PERMANOVA suggested that the fish species composition significantly differed between the depths (surface vs. bottom, F_1, 225_ = 9.21; R^2^ = 0.039; *p* < 0.001, Fig. 3), and the seasons (hypoxia vs. normoxia, F_1, 225_ = 5.37; R^2^ = 0.022; *p* < 0.001). Furthermore, the interactive term between the depth and season was also significant (F_1, 225_ = 2.58; R^2^ = 0.011; *p* < 0.001), suggesting that the degree of surface-bottom divergence differed between the two seasons. The degree of surface-bottom divergence was smaller in hypoxia, as the distance in the three-MDS dimensions between the centroids of the surface and bottom samples was 0.27 for hypoxia, and 0.50 for normoxia (Fig. 3).

**Figure 3.**
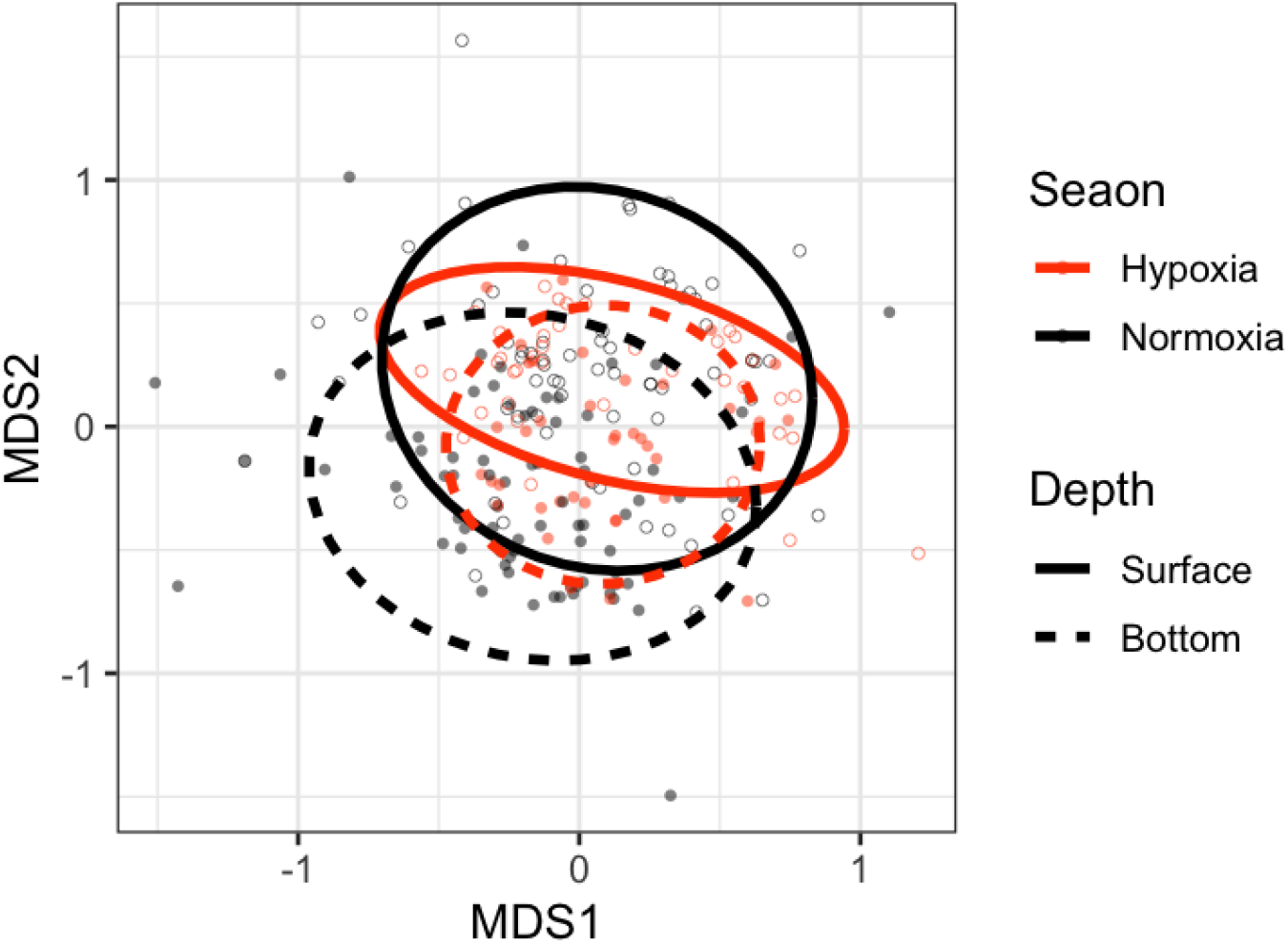
Nonmetric multidimensional scaling (NMDS) results for species composition based on Jaccard dissimilarity (dimension = 3, stress = 0.190). Each ellipse represents the 80% confidence level of the samples of each category (depth and season) in the first two MDS axes. The colour and line type of the ellipses correspond to season (orange: hypoxia, black: normoxia) and depth (solid: surface, broken: bottom), respectively. Background points similarly depict the value of the first two MDS axes of samples, with the colour and point type corresponding to season and depth (open: surface, filled: bottom), respectively.

Relative frequency of occurrence of fish species in the surface and bottom waters differed between hypoxia and normoxia. For example, *Konosirus punctatus*, the second most frequently observed species (216 out of 281 samples), occurred in the bottom water (73.3%, Fig. 4a) slightly more frequently than in the surface (72.2%) in normoxia. However, this tendency reversed in hypoxia, and it more frequently occurred in the surface (90.0%) than in the bottom (78.4%). Among the top 10% most frequently observed species (*Engraulis japonicus*, *Konosirus punctatus*, *Lateolabrax japonicus*, *Hemitrygon akajei*, *Mugil cephalus*, *Acanthopagrus schlegelii*, *Pennahia argentata*, *Callionymus valenciennei*, *Sardinella lemuru*, *Scomber japonicus*, *Sardinops sagax*, *Scomberomorus niphonius*, *Seriola quinqueradiata*, *Trachurus japonicus*, *Mustelus manazo*, *Branchiostegus japonicus* and *Trichiurus lepturus*, in order of observed frequency), such an interactive effect of depth and season on the frequency of occurrence (i.e. an increase of the relative occurrence in the surface in hypoxia) was observed for 14 out of 17 species, excepting for Japanese anchovy (*Engraulis japonicus*), Japanese sea bass (*Lateolabrax japonicus*) and Chub mackerel (*Scomber japonicus*) (Fig. 4b). The GLMM, where the occurrence of each fish species was modelled as a function of water depth, season, and their interaction with species-specific random intercept and coefficients, showed that depth and season did not consistently influence the occurrence probability (depth: coefficient = 0.228; standard error = 0.225; *p* = 0.311, season: coefficient = −0.409; standard error = 0.216, *p* = 0.058, Table 1a). On the other hand, their fixed interactive term was significant (coefficient = 0.643; standard error = 0.228; *p* = 0.005) with smaller random variance (0.328) than the other two coefficients of marginal effects (depth: 0.489, season: 0.462). Indeed, the species-specific coefficients of the interaction term estimated by random effects were positive for all the 17 species (Fig. S4). Furthermore, the likelihood test that compared the GLMM with another one, where the random effect was not modelled for the interaction term (Table 1b), showed no significant differences between the models (*χ*^2^ = 2.56, df = 1, *p* = 0.109). These results implies that the hypoxia at bottom has a negative effect on the occurrence probability consistently across species.

**Figure 4.**
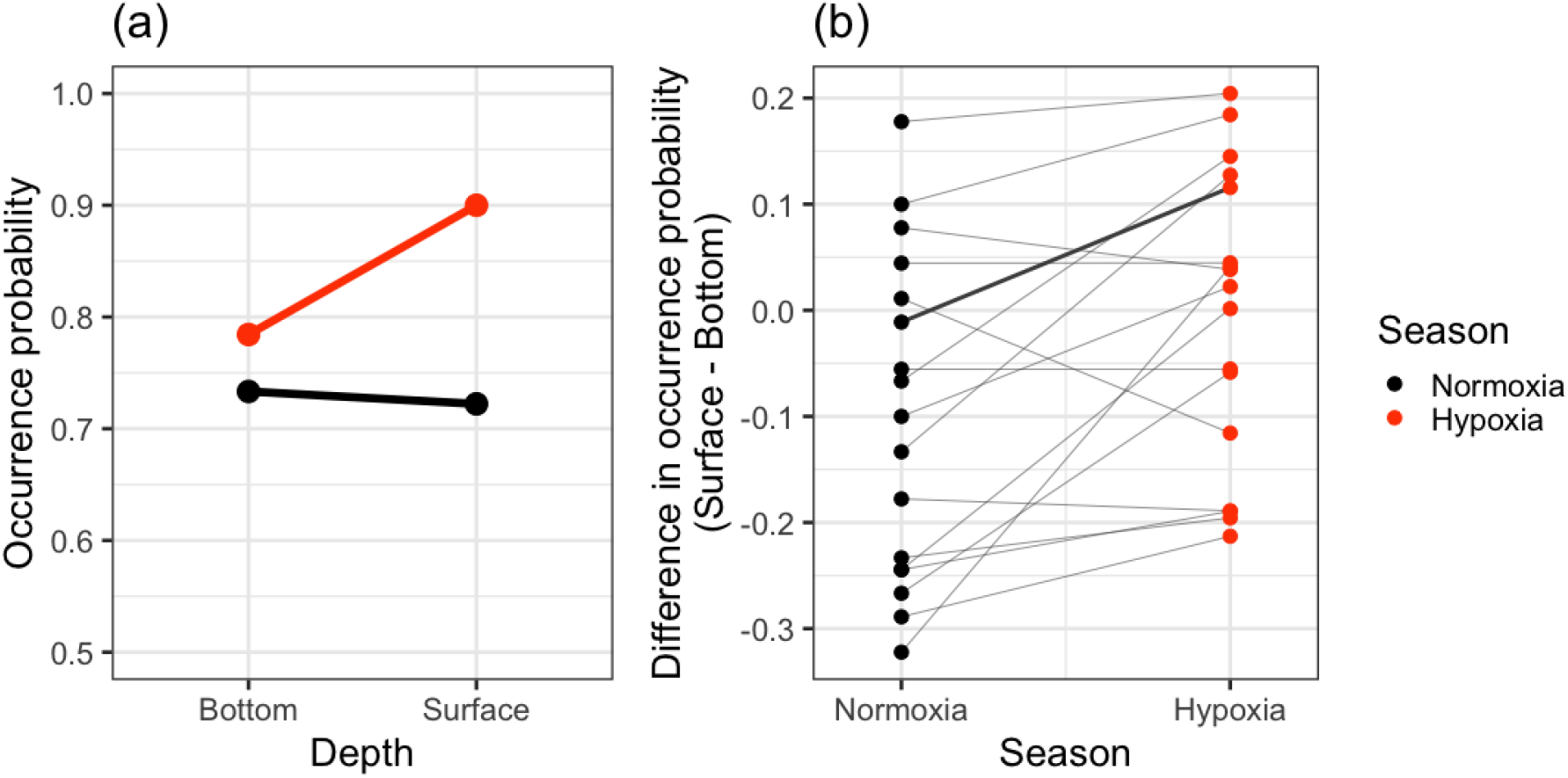
Observed relative occurrence probability in the surface and bottom waters of fish species in hypoxia and normoxia. (a) An example for *Konosirus punctatus*, the second most frequently observed species. The relative occurrence in surface compared to bottom is higher in hypoxia. (b) The difference in the occurrence probability in the surface and bottom waters in different seasons for the top 10% most frequently observed species (17 species). The points representing the same species are linked with lines, and a bold line corresponds to the example in (a). The increase of the relative occurrence in the surface in hypoxia similar to (a) was observed for 14 species.

**Table 1.**
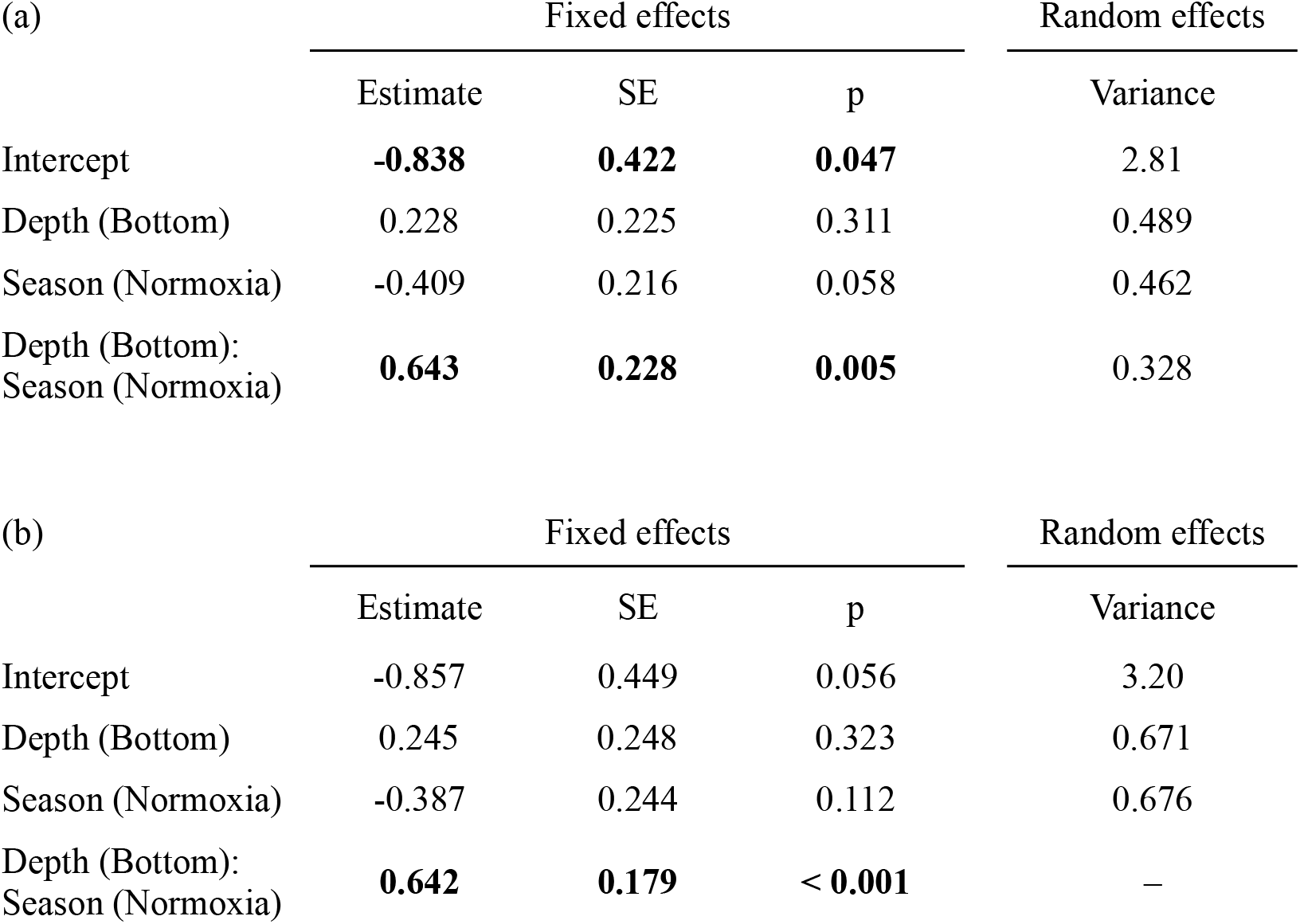
Results of the two GLMMs that modelled the presence of species as a function of water depth (surface or bottom), season (hypoxia or normoxia), and their interaction with different modelling for the random effects: (a) Species-specific random effects were modelled for the intercept, and coefficients of the three explanatory variables. (b) Species-specific random effects were modelled for the intercept and coefficients of depth and season only (not for the interaction). Estimated coefficients, standard errors (SE), *p*-values of the fixed effects, estimated variance of the random effect are shown. Significant estimates are shown with bold. The likelihood ratio test indicates that the fits of these models were not significantly different (Chi^2^ = 2.56, df = 1, *p* = 0.109).

| (a) | Fixed effects |  |  | Random effects |
| --- | --- | --- | --- | --- |
|  | Estimate | SE | p | Variance |
| Intercept | **-0.838** | **0.422** | **0.047** | 2.81 |
| Depth (Bottom) | 0.228 | 0.225 | 0.311 | 0.489 |
| Season (Normoxia) | -0.409 | 0.216 | 0.058 | 0.462 |
| Depth (Bottom):<br>Season (Normoxia) | **0.643** | **0.228** | **0.005** | 0.328 |
| (b) | Fixed effects |  |  | Random effects |
|  | Estimate | SE | p | Variance |
| Intercept | -0.857 | 0.449 | 0.056 | 3.20 |
| Depth (Bottom) | 0.245 | 0.248 | 0.323 | 0.671 |
| Season (Normoxia) | -0.387 | 0.244 | 0.112 | 0.676 |
| Depth (Bottom):<br>Season (Normoxia) | **0.642** | **0.179** | **< 0.001** | — |

## Discussion

Previous studies have argued that environmental filtering works more in abiotically more severe habitats (Chase 2007, Lepori and Malmqvist 2009, Guo et al. 2014). However, this hypothesis has rarely been tested in dynamic systems, where higher dispersal ability of species allows them to easily respond to stressful environments. In the present study system, extremely low (< 2 mg/L) DO values in the sea water were observed during summer (from June to September, Fig, 1), which likely adversely affected fish species performance and decreased the species richness in the bottom waters (Fig. 2). Based on the discussion in previous studies (Chase 2007, Lepori and Malmqvist 2009, Guo et al. 2014), the lower DO in hypoxia season was expected to strongly sort specific sets of species from the regional species pool, as tolerating such extreme oxygen conditions would require certain morphological and physiological traits (Bickler and Buck 2007, Mandic et al. 2009). Against the expectation, the results showed that filtering by DO was weak (Fig. S3) and consequently the species composition of surface and bottom waters was less distinctive (Fig. 3) during the hypoxia.

### Why weak environmental filtering during hypoxia?

These results may appear to be somewhat paradoxical, as filtering by DO was weaker during the hypoxia when the environmental conditions were more severe in the bottom and more heterogeneous between the surface and bottom waters (Fig. 1). Nevertheless, a series of evidence can explain this counterintuitive result. For example, Pihl et al. (1991) found that three demersal fish species differed in their response to bottom hypoxia in summer in Chesapeake Bay, USA. However, when the oxygen level fell below the threshold (2 mg/L), all three species similarly showed shifts in their distribution from deep to shallow waters (Pihl et al. 1991). Therefore, it is likely that, although the decrease in DO in the bottom does serve as a filter for species composition, once it exceeds a certain level of environmental severity, fish behaviors (e.g., survival, foraging, growth) will be adversely affected regardless of species identity, at which point the filtering by DO no longer plays a role in species sorting. Indeed, the GLMM in this study indicated that the random variation in the coefficient of the interaction term between water depth and hypoxia is smaller than that of depth and hypoxia (Table 1a), and the species-specific coefficients were in the same direction for all the investigated species (Fig. S4). In addition, the two GLMMs (with and without the random variation in the coefficient of the interaction term) showed similar fittings. These results collectively imply that even though a large variation in the occurrence probability in different depths and seasons exists among species (as shown by the estimated large variance of random effects, Table 1), responses to the bottom hypoxia (i.e., less frequent occurrence in the bottom waters in hypoxia) do not vary much across species. Partially supporting this conclusion that the effect of bottom hypoxia is non-species-specific, a meta-analysis revealed that effects of hypoxia on 29 fish species were not well explained by species ecological characteristics (Hrycik et al. 2017). Moreover, a recent study showed that the effect of hypoxia is variable among individuals of the same species (Steube et al. 2021), suggesting that the filtering by DO may not be precisely species-specific. Our results and other literature collectively support our second hypothesis that the filtering is weak during hypoxia because extremely low DO concentration influences the behavior of fish species in a similar way irrespective of species identity.

The observed shifts in the relative occurrence frequency from the bottom to surface waters during hypoxia can be a result of upward movements of individuals or higher mortality in the bottom (Levin 2003, Diaz and Rosenberg 2008). Here, we speculate that the former is responsible for the observed pattern. As fish are highly mobile, they can respond sharply to the environmental changes by avoiding unfavorable conditions. Therefore, when the bottom water was hypoxic in summer, they could move upward to escape from the stressful habitat (Pihl et al. 1991). This would be particularly true for free-swimming or floating species such as those frequently observed in this study. Furthermore, we observed increases in species richness per sample in the surface water during hypoxia, as well as the decreases in the bottom (Fig. S2), which supports this explanation. On the other hand, we did not obtain strong evidence for the case that higher mortality in the bottom is responsible for the observed shifts in the species occurrences. If this was the case, species richness per sample should have only decreased in the bottom and not increased in the surface, which was not in agreement with our findings (Fig. S2). Although there may indeed be higher mortality rates in the bottom water during hypoxia, it would be most influential for demersal species whose habitats are restricted to the bottom environment. However, our dataset mainly consisted of free-swimming species. Among the 17 most frequently species, only six species (*Hemitrygon akajei*, *Acanthopagrus schlegelii*, *Callionymus valenciennei*, *Seriola quinqueradiata*, *Mustelus manazo*, *Branchiostegus japonicus*) are classified as demersal based on the description in Fishbase (Froese and Pauly 2019). Though being beyond the scope of this paper, comparing the responses to bottom hypoxia for between demersal and floating fish species would be one of the fruitful investigations of future studies.

### Implications for community assembly studies

Our results have general implication in the context of community assembly. Previous studies have shown that deterministic assembly plays a more important role in stressful habitats because the severe environment filters potential colonizers (Chase 2007, Lepori and Malmqvist 2009, Guo et al. 2014). Our results are inconsistent with this in that deterministic control by environments was weaker when the environment was more severe. Nevertheless, a limited number of studies have suggested that this may be the case. Lepori and Malmqvist (2009) have found that the deterministic control is strongest at the intermediate level of disturbance. They agreed with the previous discussion that more frequent disturbance intensified species sorting by selecting disturbance-tolerant species, but they concluded that the disturbance that was too severe induced extinction of individuals irrespective of species traits (Lepori and Malmqvist 2009), which would hamper the environmental filtering. Similarly, Kim (2019) pointed to a possibility that species living in extremely severe environments share similar tolerance traits. If this was the case, environmental filtering may no longer play a role in such “neutral” communities (Kim 2019). Our findings add to this series of evidence, by showing that weaker environmental filtering was acting in more stressful environments.

Furthermore, the results showing weaker filtering in summer when the surface-bottom gradient of DO was more distinct (Fig. 1) challenge the pervasive view that stronger filtering is expected when the environmental gradient was more heterogeneous (Kraft et al. 2015). Based on our results, we speculate that although increasing environmental heterogeneity may indeed lead to stronger filtering, once the environmental gradient becomes wide enough to encompass extremely severe conditions, the filtering would become weak.

### Caveats and utility of eDNA analyses

Although the ability of eDNA to detect species from environmental samples has been established (Thomsen et al. 2012; Sigsgaard et al. 2017; Yamamoto et al. 2017), this technique has rarely been applied to the evaluation of community assembly processes. In this study, the eDNA analysis successfully reveals the seasonal shifts in the strength of environmental filtering according to hypoxia, which illustrates its capability to be applied to community assembly studies. However, the dataset we used contained only presence–absence information, which would have missed some important information. Estimating species abundance data from eDNA samples is challenging at the present time as the correlation between eDNA concentration and abundance of target fish species has not been established under natural conditions (Yates et al. 2019). To avoid potential biases, we did not use the read number of DNA as an abundance index. Moreover, the detection of eDNA is influenced not only by the presence of the species but also by the degradation and accumulation of the eDNA itself (“ecology of eDNA” sensu Barnes and Turner 2016). For example, it is known that the degradation of eDNA after release into water is faster at higher temperatures (Tsuji et al. 2017, Jo et al. 2019), and it has been assumed that eDNA accumulates in surface waters (Eichmiller et al. 2014, Moyer et al. 2014). We consider that these biases are not the case here or at least do not change our main finding, because species richness in surface water was high in summer where the degradation is expected to be faster (Fig. S2), and the richness was generally higher in the bottom where the accumulation is expected to be reduced (Fig. S2).

## Conclusion

In ecosystems, species assemblages are determined by abiotic environmental conditions, but to a variable extent. We found that the extremely severe environment, that is the seasonal bottom hypoxia in coastal ecosystems, made the filtering weaker likely because its adverse effect was non-species-specific. This result may seem to be contradictory to the current pervasive view that deterministic control by abiotic conditions is stronger in more stressful habitats. However, we speculate that these two contrasting conclusions are compatible: While the increasing environmental severity surely strengthens the filtering on species composition by sorting out vulnerable species, once it exceeds a certain level, the filtering would no longer work because environments that are too severe harm all the species in a similar way. Therefore, the deterministic control is expected to be strongest at the intermediate level of environmental severity, a hypothesis to be further confirmed in the future studies. Furthermore, by applying the eDNA analyses for detecting community assembly processes for the first time to our best knowledge, this study highlights the potential applications of this promising technique across a wide range of disciplines of ecology.

## References

Anderson, M. J. M. 2001. A new method for non-parametric multivariate analysis of variance. Austral Ecology 26:32–46.

Barnes, M. A., and C. R. Turner. 2016. The ecology of environmental DNA and implications for conservation genetics. Conservation Genetics 17:1–17.

Bickler, P. E., and L. T. Buck. 2007. Hypoxia tolerance in reptiles, amphibians, and fishes: Life with variable oxygen availability. Annual Review of Physiology.

Bohmann, K., A. Evans, M. T. P. Gilbert, G. R. Carvalho, S. Creer, M. Knapp, D. W. Yu, et al. 2014. Environmental DNA for wildlife biology and biodiversity monitoring. Trends in Ecology and Evolution 29:358–367.

Breitburg, D., L. A. Levin, A. Oschlies, M. Grégoire, F. P. Chavez, D. J. Conley, V. Garçon, et al. 2018. Declining oxygen in the global ocean and coastal waters. Science 359.

Brooks, M. E., K. Kristensen, K. J. van Benthem, A. Magnusson, C. W. Berg, A. Nielsen, H. J. Skaug, et al. 2017. glmmTMB Balances Speed and Flexibility Among Packages for Zero-inflated Generalized Linear Mixed Modeling. The R Journal, 9(2), 378–400. https://journal.r-project.org/archive/2017/RJ-2017-066/index.html.

Chase, J. M. 2007. Drought mediates the importance of stochastic community assembly. Proceedings of the National Academy of Sciences of the United States of America 104:17430–17434.

Chase, J. M., and J. A. Myers. 2011. Disentangling the importance of ecological niches from stochastic processes across scales. Philosophical Transactions of the Royal Society B: Biological Sciences 366:2351–2363.

Cottenie, K. 2005. Integrating environmental and spatial processes in ecological community dynamics. Ecology Letters 8:1175–1182.

Deiner, K., H. M. Bik, E. Mächler, M. Seymour, A. Lacoursière-Roussel, F. Altermatt, S. Creer, et al. 2017. Environmental DNA metabarcoding: Transforming how we survey animal and plant communities. Molecular Ecology 26:5872–5895.

Diaz, R. J., and R. Rosenberg. 1995. Marine benthic hypoxia: A review of its ecological effects and the behavioural response of benthic macrofauna MARINE. Oceanography and Morine Biology: an Annual Review 33:245–303.

Diaz, R. J., and R. Rosenberg. 2008. Spreading Dead Zones and Consequences for Marine Ecosystems. Science 321:926–929.

Eichmiller, J. J., P. G. Bajer, and P. W. Sorensen. 2014. The relationship between the distribution of common carp and their environmental DNA in a small lake. PLoS ONE 9:1–8.

Froese, R., and D. Pauly. (Eds). 2019. FishBase. World Wide Web electronic publication. www.fishbase.org, version (12/2019).

Grinnell, J. 1917. The Niche-Relationships of the California Thrasher. The Auk 34:427–433.

Guo, H., K. Wieski, Z. Lan, and S. C. Pennings. 2014. Relative influence of deterministic processes on structuring marsh plant communities varies across an abiotic gradient. Oikos 123:173–178.

HilleRisLambers, J., P. B. Adler, W. S. Harpole, J. M. Levine, and M. M. Mayfield. 2012. Rethinking Community Assembly through the Lens of Coexistence Theory. Annual Review of Ecology, Evolution, and Systematics 43:227–248.

Hongo, Y., S. Nishijima, Y. Kanamori, S. Sawayama, K. Yokouchi, N. Kanda, S. Oori, et al. 2021. Fish environmental DNA in Tokyo Bay: A feasibility study on the availability of environmental DNA for fisheries. Regional Studies in Marine Science 47:101950.

Hrycik, A. R., L. Z. Almeida, and T. O. Höök. 2017. Sub-lethal effects on fish provide insight into a biologically-relevant threshold of hypoxia. Oikos 126:307–317.

Hubbell, S. P. 2001. The Unified Neutral Theory of Biodiversity and Biogeography.

Hutchinson, G. E. 1959. Homage to Santa Rosalia or Why Are There So Many Kinds of Animals? The American Naturalist 93:145–159.

Jo, T., H. Murakami, S. Yamamoto, R. Masuda, and T. Minamoto. 2019. Effect of water temperature and fish biomass on environmental DNA shedding, degradation, and size distribution. Ecology and Evolution 9:1135–1146.

Kim, D. 2019. Selection of scale can reverse the importance of stochastic controls on community assembly. Physical Geography 40:111–126.

Kodama, K., and T. Horiguchi. 2011. Effects of hypoxia on benthic organisms in Tokyo Bay, Japan: A review. Marine Pollution Bulletin 63:215–220.

Kouno, H., H. Kanou, and T. Yokoo. (Eds.) 2011. A Photographic Guide to the Fishes in Tokyo Bay. Tokyo, Heibon-sha. [in Japanese]

Kraft, N. J. B., P. B. Adler, O. Godoy, E. C. James, S. Fuller, and J. M. Levine. 2015. Community assembly, coexistence and the environmental filtering metaphor. Functional Ecology 29:592–599.

Legendre, P., and M. J. Andersson. 1999. Distance-based redundancy analysis: Testing multispecies responses in multifactorial ecological experiments. Ecological Monographs 69:1–24.

Lepori, F., and B. Malmqvist. 2009. Deterministic control on community assembly peaks at intermediate levels of disturbance. Oikos 118:471–479.

Levin, L. A. 2003. OXYGEN MINIMUM ZONE BENTHOS: ADAPTATION AND COMMUNITY RESPONSE TO HYPOXIA. Oceanography and Marine Biology: an Annual Review 41:1–45.

Mandic, M., A. E. Todgham, and J. G. Richards. 2009. Mechanisms and evolution of hypoxia tolerance in fish. Proceedings of the Royal Society B: Biological Sciences 276:735–744.

Miya, M., Y. Sato, T. Fukunaga, T. Sado, J. Y. Poulsen, K. Sato, T. Minamoto, et al. 2015. MiFish, a set of universal PCR primers for metabarcoding environmental DNA from fishes: Detection of more than 230 subtropical marine species. Royal Society Open Science 2.

Moyer, G. R., E. Díaz-Ferguson, J. E. Hill, and C. Shea. 2014. Assessing environmental DNA detection in controlled lentic systems. PLoS ONE 9.

Murakami, H., S. Yoon, A. Kasai, T. Minamoto, S. Yamamoto, M. K. Sakata, T. Horiuchi, et al. 2019. Dispersion and degradation of environmental DNA from caged fish in a marine environment. Fisheries Science 85:327–337.

Nakabo, T. (Ed.) 2002. Fishes of Japan with Pictorial Keys to the Species. Kanagawa, Tokai University Press. [in Japanese]

Oksanen, J., F. G. Blanchet, M. Friendly, R. Kindt, P. Legendre, D. McGlinn, P. R. Minchin, R. B. O’Hara, G. L. Simpson, P. Solymos, et al. 2019. vegan: Community Ecology Package. R package version 2.5-6. https://CRAN.R-project.org/package=vegan

Peres-Neto, P. R., P. Legendre, S. Dray, and D. Borcard. 2006. Variation partitioning of species data matrices: Estimation and comparison of fractions. Ecology 87:2614–2625.

Pihl, L., S. P. Baden, and R. J. Diaz. 1991. Effects of periodic hypoxia on distribution of demersal fish and crustaceans. Marine Biology 108:349–360.

Port, J. A., J. L. O’Donnell, O. C. Romero-Maraccini, P. R. Leary, S. Y. Litvin, K. J. Nickols, K. M. Yamahara, et al. 2016. Assessing vertebrate biodiversity in a kelp forest ecosystem using environmental DNA. Molecular Ecology 25:527–541.

R Core Team. 2020. R: A language and environment for statistical computing. R Foundation for Statistical Computing, Vienna, Austria. URL https://www.R-project.org/

Sale, P. F. 1978. Coexistence of coral reef fishes - a lottery for living space. Environmental Biology of Fishes 3:85–102.

Sato, Y., M. Miya, T. Fukunaga, T. Sado, and W. Iwasaki. 2018. MitoFish and mifish pipeline: A mitochondrial genome database of fish with an analysis pipeline for environmental DNA metabarcoding. Molecular Biology and Evolution 35:1553–1555.

Sigsgaard, E. E., I. B. Nielsen, H. Carl, M. A. Krag, S. W. Knudsen, Y. Xing, T. H. Holm-Hansen, et al. 2017. Seawater environmental DNA reflects seasonality of a coastal fish community. Marine Biology 164:1–15.

Steube, T. R., M. E. Altenritter, and B. D. Walther. 2021. Distributive stress: individually variable responses to hypoxia expand trophic niches in fish. Ecology 102.

Thomsen, P. F., J. Kielgast, L. L. Iversen, P. R. Møller, M. Rasmussen, and E. Willerslev. 2012. Detection of a Diverse Marine Fish Fauna Using Environmental DNA from Seawater Samples. PLoS ONE 7:1–9.

Tilman, D., S. S. Kilham, and P. Kilham. 1982. Phytoplankton community ecology: the role of limiting nutrients. Annual review of ecology and systematics. Volume 13 13:349–372.

Tsuji, S., M. Ushio, S. Sakurai, T. Minamoto, and H. Yamanaka. 2017. Water temperature-dependent degradation of environmental DNA and its relation to bacterial abundance. PLoS ONE 12:1–13.

Wickham H 2016. ggplot2: Elegant Graphics for Data Analysis. Springer-Verlag New York. ISBN 978-3-319-24277-4, https://ggplot2.tidyverse.org.

Yamamoto, S., R. Masuda, Y. Sato, T. Sado, H. Araki, M. Kondoh, T. Minamoto, et al. 2017. Environmental DNA metabarcoding reveals local fish communities in a species-rich coastal sea. Scientific Reports 7:1–12.

Yamamoto, S., K. Minami, K. Fukaya, K. Takahashi, H. Sawada, H. Murakami, S. Tsuji, et al. 2016. Environmental DNA as a “snapshot” of fish distribution: A case study of Japanese jack mackerel in Maizuru Bay, Sea of Japan. PLoS ONE 11:1–18.

Yasui, S., J. Kanda, T. Usui, and H. Ogawa. 2016. Seasonal variations of dissolved organic matter and nutrients in sediment pore water in the inner part of Tokyo Bay. Journal of Oceanography 72:851–866.

Yates, M. C., D. J. Fraser, and A. M. Derry. 2019. Meta□analysis supports further refinement of eDNA for monitoring aquatic species□specific abundance in nature. Environmental DNA 1:5–13.

