## Supplemental materials for "Environmental filtering in marine fish communities is weakened during severe hypoxia season"

**Supplementary materials**

Figure S1.	Map of the 14 sampling sites in Tokyo Bay, Japan.

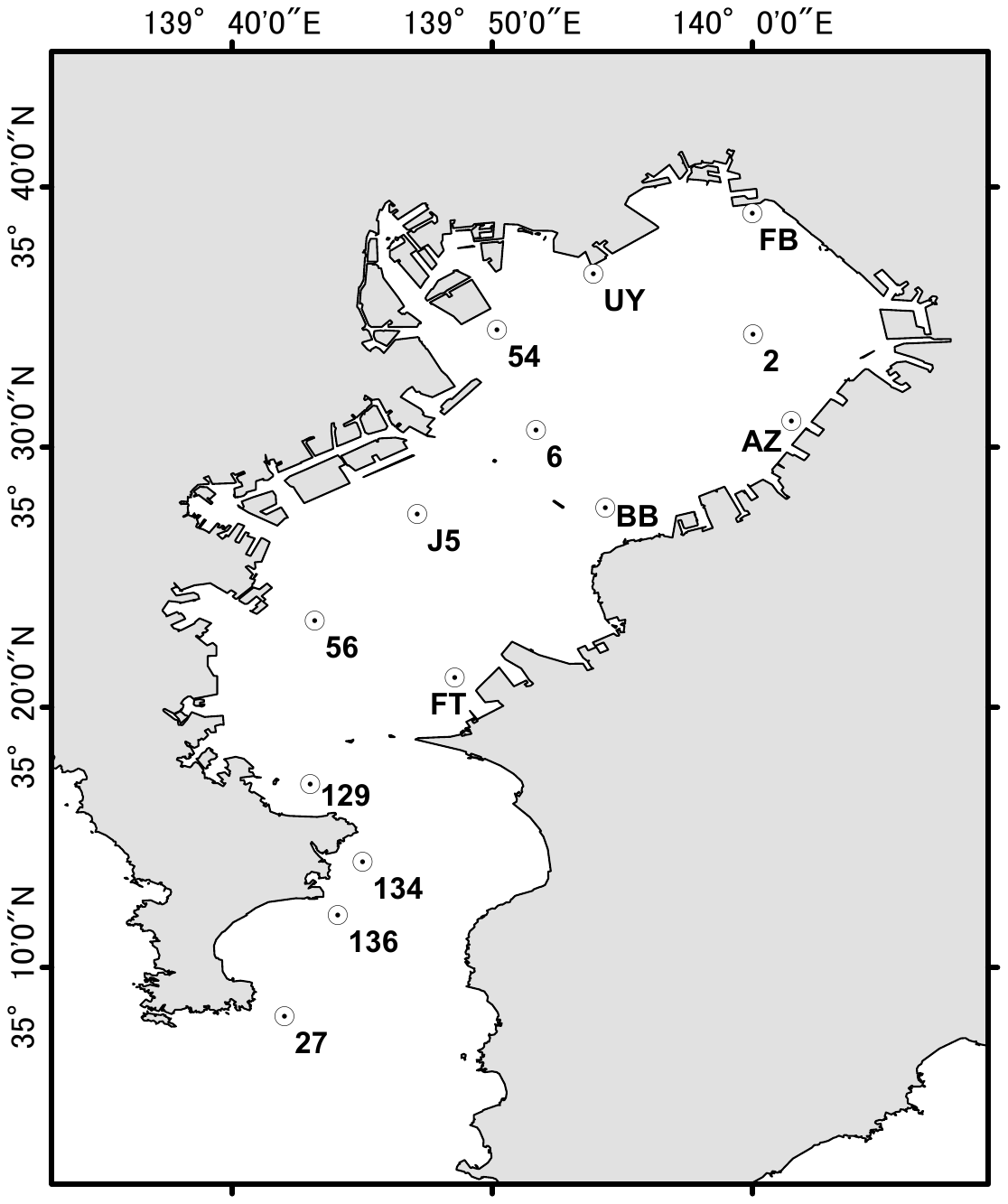

Figure S2.	Number of species per samples of different months and depths. The period of hypoxia (from June to September) is bordered with red.

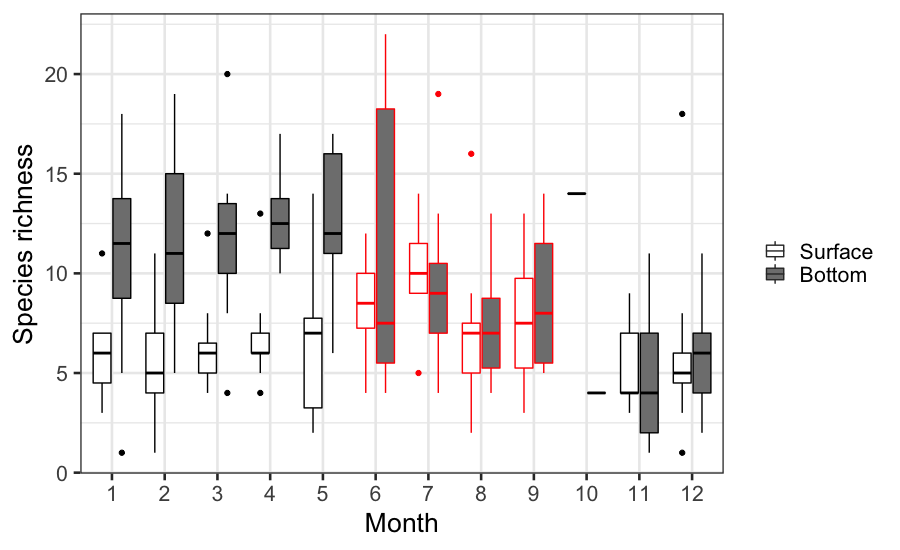

Figure S3.	Difference in the variance of fish species composition explained by dissolved oxygen. R^2^ values were obtained from distance-based redundancy analyses that were separately conducted for each month except for October.

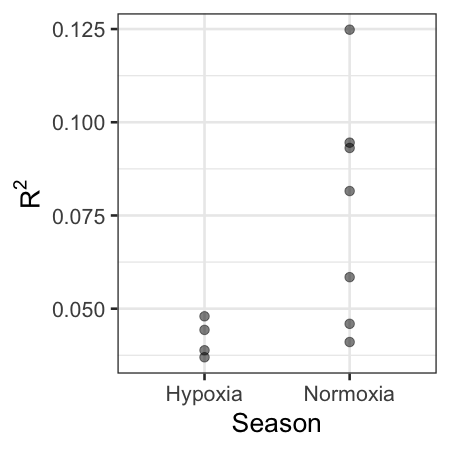

Figure S4.	Estimated species-specific coefficients of the interactive term of the depth and season of the GLMM. The means and confidence intervals (1.96 standard error　ranges) are shown for each species. The species ID number represents the order of occurrence, with species 1 being most frequently observed (species names are shown in the main text with their order of observed frequency).

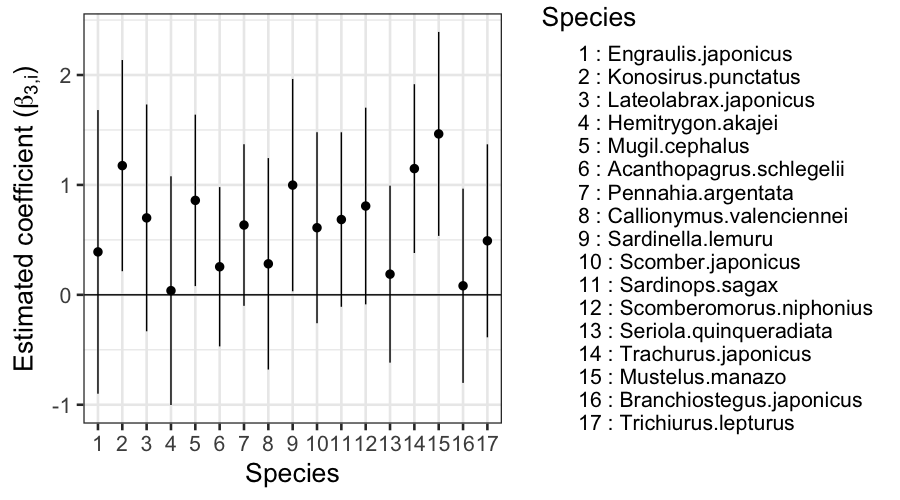

Table S1.	List of the fish species detected by our sequencing analyses. The third and fourth columns represent their species name found in the Mitofish database, and that matched in Fishbase, respectively. Therefore, NAs in the fourth column indicate filtering by our first criterion (*Material and methods* in the main text). The third column contains information on our second criterion that judged whether the species habituates in Tokyo Bay.

| **Order** | **Family** | **Species name**  **(MitoFish)** | **Species name**  **(Fishbase)** | **Distribution** |
| --- | --- | --- | --- | --- |
| Myxiniformes | Myxinidae | Eptatretus burgeri | Eptatretus burgeri | In |
| Lamniformes | Alopiidae | Alopias vulpinus | Alopias vulpinus | In |
| Carcharhiniformes | Carcharhinidae | Carcharhinus  limbatus | Carcharhinus  limbatus | Out |
|  | Sphyrnidae | Sphyrna zygaena | Sphyrna zygaena | In |
|  | Triakidae | Mustelus griseus | Mustelus griseus | In |
|  |  | Mustelus manazo | Mustelus manazo | In |
|  |  | Triakis scyllium | Triakis scyllium | In |
| Squatiniformes | Squatinidae | Squatina japonica | Squatina japonica | In |
| Torpediniformes | Narkidae | Narke japonica | Narke japonica | In |
| Rajiformes | Arhynchobatidae | Bathyraja smirnovi | Bathyraja smirnovi | Out |
|  | Rajidae | Dipturus  kwangtungensis | Dipturus  kwangtungensis | In |
|  |  | Okamejei  acutispina | Okamejei  acutispina | In |
| Myliobatiformes | Dasyatidae | Bathytoshia  brevicaudata | Bathytoshia  brevicaudata | Out |
|  |  | Hemitrygon akajei | Hemitrygon akajei | In |
|  |  | Hemitrygon  izuensis | Hemitrygon  izuensis | In |
|  |  | Dasyatis say | Hypanus say | Out |
|  | Gymnuridae | Gymnura japonica | Gymnura japonica | In |
|  | Myliobatidae | Myliobatis tobijei | Myliobatis tobijei | In |
|  | Rhinopteridae | Rhinoptera  bonasus | Rhinoptera  bonasus | Out |
|  | Urolophidae | Urolophus  aurantiacus | Urolophus  aurantiacus | In |
| Elopiformes | Elopidae | Elops hawaiensis | Elops hawaiensis | In |
| Anguilliformes | Congridae | Conger myriaster | Conger myriaster | In |
|  | Muraenesocidae | Muraenesox  cinereus | Muraenesox  cinereus | In |
|  | Muraenidae | Gymnothorax  kidako | Gymnothorax  kidako | In |
|  | Nemichthyidae | Nemichthys  scolopaceus | Nemichthys  scolopaceus | In |
|  | Ophichthidae | Brachysomophis  crocodilinus | Brachysomophis  crocodilinus | Out |
|  |  | Ophichthus  zophistius | Ophichthus  altipennis | In |
|  |  | Ophisurus  macrorhynchos | Ophisurus  macrorhynchos | In |
|  |  | Scolecenchelys  borealis | Scolecenchelys  aoki | Out |
| Clupeiformes | Clupeidae | Konosirus  punctatus | Konosirus  punctatus | In |
|  |  | Sardinella lemuru | Sardinella lemuru | In |
|  |  | Sardinops  melanostictus | Sardinops sagax | In |
|  | Dussumieriidae | Etrumeus teres | Etrumeus sadina | In |
|  | Engraulidae | Engraulis  japonicus | Engraulis  japonicus | In |
| Cypriniformes | Cyprinidae | Carassius auratus  grandoculis | Carassius auratus | Out |
|  |  | Carassius auratus x  Megalobrama  amblycephala  pentaploid hybrid | NA | NA |
|  |  | Carassius sp.  'Ginbuna' | NA | NA |
|  |  | Cyprinus carpio | Cyprinus carpio | Out |
|  |  | Cyprinus carpio  'Ying hybrid' | NA | NA |
|  |  | Hemibarbus barbus | Hemibarbus labeo | Out |
|  |  | Hypophthalmichthys  molitrix | Hypophthalmichthys  molitrix | Out |
|  |  | Hypophthalmichthys  nobilis x  Hypophthalmichthys  molitrix | NA | NA |
|  |  | Phoxinus phoxinus  tumensis | NA | NA |
|  |  | Phoxinus  steindachneri | Phoxinus  steindachneri | Out |
|  |  | Pseudorasbora  parva | Pseudorasbora  parva | Out |
|  |  | Tribolodon  brandtii | Tribolodon  brandtii | In |
|  |  | Tribolodon  hakonensis | Tribolodon  hakonensis | In |
|  | Nemacheilidae | Barbatula  barbatula | Barbatula  barbatula | Out |
|  |  | Misgurnus  anguillicaudatus | Misgurnus  anguillicaudatus | Out |
|  |  | Misgurnus  anguillicaudatus x  Paramisgurnus  dabryanus | NA | NA |
| Siluriformes | Amblycipitidae | Liobagrus reinii | Liobagrus reinii | Out |
|  | Plotosidae | Plotosus japonicus | Plotosus japonicus | In |
| Osmeriformes | Plecoglossidae | Plecoglossus  altivelis | Plecoglossus  altivelis | In |
| Salmoniformes | Salmonidae | Oncorhynchus keta | Oncorhynchus keta | Out |
|  |  | Oncorhynchus  mykiss | Oncorhynchus  mykiss | In |
|  |  | Oncorhynchus  nerka | Oncorhynchus  nerka | Out |
| Stomiiformes | Gonostomatidae | Sigmops gracilis | Sigmops gracilis | In |
|  | Phosichthyidae | Woodsia  nonsuchae | Woodsia  nonsuchae | Out |
|  | Sternoptychidae | Maurolicus  muelleri | Maurolicus  muelleri | In |
| Aulopiformes | Aulopidae | Hime japonica | Hime japonica | In |
|  | Synodontidae | Saurida microlepis | Saurida microlepis | Out |
| Myctophiformes | Myctophidae | Benthosema  pterotum | Benthosema  pterotum | In |
|  |  | Diaphus  suborbitalis | Diaphus  suborbitalis | In |
|  |  | Hygophum  reinhardtii | Hygophum  reinhardtii | In |
| Gadiformes | Gadidae | Gadus  chalcogrammus | Gadus  chalcogrammus | In |
|  | Macrouridae | Coryphaenoides  marginatus | Coryphaenoides  marginatus | In |
| Lophiiformes | Lophiidae | Lophius litulon | Lophius litulon | In |
| Atheriniformes | Atherinidae | Hypoatherina  valenciennei | Hypoatherina  valenciennei | In |
| Cyprinodontiformes | Poeciliidae | Gambusia affinis | Gambusia affinis | Out |
| Beloniformes | Belonidae | Hyporhamphus  sajori | Hyporhamphus  sajori | In |
|  |  | Strongylura  anastomella | Strongylura  anastomella | In |
|  |  | Tylosurus  crocodilus  crocodilus | Tylosurus  crocodilus | In |
|  | Exocoetidae | Hirundichthys  rondeletii | Hirundichthys  rondeletii | In |
|  | Scomberesocidae | Cololabis saira | Cololabis saira | In |
| Beryciformes | Berycidae | Beryx splendens | Beryx splendens | In |
| Syngnathiformes | Syngnathidae | Solegnathus  hardwickii | Solegnathus  hardwickii | Out |
| Scorpaeniformes | Cottidae | Cottus asper | Cottus asper | Out |
|  |  | Cottus  hangiongensis | Cottus  hangiongensis | Out |
|  |  | Pseudoblennius  percoides | Pseudoblennius  percoides | In |
|  | Hexagrammidae | Hexagrammos  otakii | Hexagrammos  otakii | In |
|  |  | Pleurogrammus  monopterygius | Pleurogrammus  monopterygius | Out |
|  | Platycephalidae | Cociella crocodila | Cociella crocodilus | In |
|  |  | Platycephalus sp.  MAGOCHI | NA | NA |
|  |  | Suggrundus  meerdervoortii | Suggrundus  meerdervoortii | In |
|  | Sebastidae | Scorpaenodes  evides | Scorpaenodes  evides | In |
|  |  | Sebastes aleutianus | Sebastes aleutianus | Out |
|  |  | Sebastes  nigrocinctus | Sebastes  nigrocinctus | Out |
|  |  | Sebastiscus  albofasciatus | Sebastiscus  albofasciatus | In |
|  |  | Sebastiscus  marmoratus | Sebastiscus  marmoratus | In |
|  | Synanceiidae | Inimicus japonicus | Inimicus japonicus | In |
|  | Tetrarogidae | Paracentropogon  rubripinnis | Paracentropogon  rubripinnis | In |
|  | Triglidae | Lepidotrigla  abyssalis | Lepidotrigla  abyssalis | In |
|  |  | Lepidotrigla  kishinouyi | Lepidotrigla  kishinouyi | In |
|  |  | Lepidotrigla  microptera | Lepidotrigla  microptera | In |
|  |  | Lepidotrigla sp.  NMMBP 1282 | NA | NA |
|  |  | Eutrigla gurnardus | Eutrigla gurnardus | Out |
| Perciformes | Acropomatidae | Malakichthys  griseus | Malakichthys  griseus | In |
|  | Apogonidae | Jaydia lineata | Jaydia lineata | In |
|  |  | Apogon  semilineatus | Ostorhinchus  semilineatus | In |
|  | Blenniidae | Parablennius  yatabei | Parablennius  yatabei | In |
|  |  | Omobranchus  punctatus | Omobranchus  punctatus | In |
|  | Callionymidae | Callionymus  beniteguri | Callionymus  beniteguri | In |
|  |  | Callionymus  enneactis | Callionymus  enneactis | In |
|  |  | Callionymus  japonicus | Callionymus  japonicus | In |
|  |  | Callionymus  valenciennei | Callionymus  valenciennei | In |
|  | Carangidae | Decapterus  macarellus | Decapterus  macarellus | In |
|  |  | Decapterus  maruadsi | Decapterus  maruadsi | In |
|  |  | Trachurus  japonicus | Trachurus  japonicus | In |
|  |  | Seriola dumerili | Seriola dumerili | In |
|  |  | Seriola  quinqueradiata | Seriola  quinqueradiata | In |
|  | Centrarchidae | Lepomis  macrochirus | Lepomis  macrochirus | Out |
|  | Centrolophidae | Hyperoglyphe  japonica | Hyperoglyphe  japonica | In |
|  |  | Psenopsis anomala | Psenopsis anomala | In |
|  | Chaenopsidae | Neoclinus nudus | Neoclinus nudus | Out |
|  | Chaetodontidae | Chaetodon  modestus | Roa modesta | In |
|  | Cheilodactylidae | Cheilodactylus  zonatus | Cheilodactylus  zonatus | In |
|  | Eleotridae | Eleotris  oxycephala | Eleotris  oxycephala | Out |
|  | Embiotocidae | Ditrema viride | Ditrema viride | In |
|  |  | Neoditrema  ransonnetii | Neoditrema  ransonnetii | In |
|  | Gempylidae | Gempylus serpens | Gempylus serpens | In |
|  |  | Promethichthys  prometheus | Promethichthys  prometheus | In |
|  | Girellidae | Girella punctata | Girella punctata | In |
|  | Gobiidae | Acanthogobius  flavimanus | Acanthogobius  flavimanus | In |
|  |  | Acentrogobius  virgatulus | NA | NA |
|  |  | Amblychaeturichthys  hexanema | Amblychaeturichthys  hexanema | In |
|  |  | Amblychaeturichthys  sciistius | Amblychaeturichthys  sciistius | In |
|  |  | Chaenogobius  gulosus | Chaenogobius  gulosus | In |
|  |  | Favonigobius  gymnauchen | Favonigobius  gymnauchen | In |
|  |  | Gymnogobius  cylindricus | Gymnogobius  cylindricus | Out |
|  |  | Gymnogobius  heptacanthus | Gymnogobius  heptacanthus | In |
|  |  | Mugilogobius abei | Mugilogobius abei | In |
|  |  | Myersina filifer | Myersina filifer | In |
|  |  | Pterogobius virgo | Pterogobius virgo | In |
|  |  | Pterogobius  zacalles | Pterogobius  zacalles | In |
|  |  | Rhinogobius  brunneus | Rhinogobius  brunneus | In |
|  |  | Sagamia  geneionema | Sagamia  geneionema | In |
|  |  | Tridentiger  bifasciatus | Tridentiger  bifasciatus | In |
|  |  | Tridentiger  kuroiwae | Tridentiger  kuroiwae | Out |
|  |  | Tridentiger  obscurus | Tridentiger  obscurus | In |
|  |  | Tridentiger  trigonocephalus | Tridentiger  trigonocephalus | In |
|  | Haemulidae | Parapristipoma  trilineatum | Parapristipoma  trilineatum | In |
|  |  | Plectorhinchus  cinctus | Plectorhinchus  cinctus | In |
|  | Istiophoridae | Istiophorus  platypterus | Istiophorus  platypterus | In |
|  | Labridae | Halichoeres  tenuispinis | Halichoeres  tenuispinis | In |
|  |  | Parajulis  poecilepterus | Parajulis  poecilepterus | In |
|  |  | Pseudolabrus  sieboldi | Pseudolabrus  sieboldi | In |
|  |  | Pteragogus  aurigarius | Pteragogus  aurigarius | In |
|  |  | Semicossyphus  reticulatus | Semicossyphus  reticulatus | In |
|  | Lateolabracidae | Lateolabrax  japonicus | Lateolabrax  japonicus | In |
|  | Leiognathidae | Equulites rivulatus | Equulites rivulatus | In |
|  |  | Nuchequula  nuchalis | Nuchequula  nuchalis | In |
|  | Lobotidae | Lobotes  surinamensis | Lobotes  surinamensis | In |
|  | Malacanthidae | Branchiostegus  japonicus | Branchiostegus  japonicus | In |
|  | Microdesmidae | Parioglossus dotui | Parioglossus dotui | In |
|  | Nemipteridae | Nemipterus  bathybius | Nemipterus  bathybius | In |
|  | Neozoarcidae | Zoarchias glaber | Zoarchias glaber | In |
|  | Oplegnathidae | Oplegnathus  fasciatus | Oplegnathus  fasciatus | In |
|  | Pholidae | Pholis nebulosa | Pholis nebulosa | In |
|  | Pinguipedidae | Parapercis  diplospilus | Parapercis  diplospilus | Out |
|  |  | Parapercis  multifasciata | Parapercis  multifasciata | In |
|  |  | Parapercis  pulchella | Parapercis  pulchella | In |
|  |  | Parapercis  sexfasciata | Parapercis  sexfasciata | In |
|  |  | Parapercis snyderi | Parapercis snyderi | In |
|  | Pomacentridae | Chromis fumea | Chromis fumea | In |
|  |  | Chromis notata  notata | Chromis notata | In |
|  | Sciaenidae | Nibea mitsukurii | Nibea mitsukurii | In |
|  |  | Pennahia argentata | Pennahia argentata | In |
|  | Scombridae | Auxis rochei | Auxis rochei | In |
|  |  | Katsuwonus  pelamis | Katsuwonus  pelamis | In |
|  |  | Scomber japonicus | Scomber japonicus | In |
|  |  | Scomberomorus  niphonius | Scomberomorus  niphonius | In |
|  |  | Thunnus alalunga | Thunnus alalunga | In |
|  |  | Thunnus albacares | Thunnus albacares | In |
|  |  | Thunnus maccoyii | Thunnus maccoyii | Out |
|  |  | Thunnus thynnus  thynnus | Thunnus thynnus | Out |
|  | Scombropidae | Scombrops boops | Scombrops boops | In |
|  |  | Scombrops gilberti | Scombrops gilberti | In |
|  | Serranidae | Chelidoperca  hirundinacea | Chelidoperca  hirundinacea | In |
|  |  | Chelidoperca  pleurospilus | Chelidoperca  pleurospilus | In |
|  |  | Epinephelus  fasciatus | Epinephelus  fasciatus | In |
|  |  | Niphon spinosus | Niphon spinosus | In |
|  |  | Pseudanthias  rubrolineatus | Pseudanthias  rubrolineatus | Out |
|  |  | Sacura  margaritacea | Sacura  margaritacea | In |
|  | Siganidae | Siganus  canaliculatus | Siganus  canaliculatus | Out |
|  |  | Siganus fuscescens | Siganus fuscescens | In |
|  | Sillaginidae | Sillago japonica | Sillago japonica | In |
|  | Sparidae | Acanthopagrus  latus | Acanthopagrus  latus | In |
|  |  | Acanthopagrus  schlegelii | Acanthopagrus  schlegelii | In |
|  |  | Acanthopagrus  schlegelii x Pagrus major | NA | NA |
|  |  | Dentex  hypselosomus | Dentex  hypselosomus | In |
|  |  | Dentex tumifrons | Dentex tumifrons | In |
|  | Sphyraenidae | Sphyraena  japonica | Sphyraena  japonica | In |
|  |  | Sphyraena pinguis | Sphyraena pinguis | In |
|  | Stichaeidae | Dictyosoma  burgeri | Dictyosoma  burgeri | In |
|  |  | Dictyosoma  rubrimaculatum | Dictyosoma  rubrimaculatum | In |
|  | Terapontidae | Rhynchopelates  oxyrhynchus | Rhynchopelates  oxyrhynchus | In |
|  | Trichiuridae | Trichiurus lepturus | Trichiurus lepturus | In |
|  | Xiphiidae | Xiphias gladius | Xiphias gladius | In |
| Mugiliformes | Mugilidae | Mugil cephalus | Mugil cephalus | In |
| Pleuronectiformes | Cynoglossidae | Cynoglossus  interruptus | Cynoglossus  interruptus | In |
|  |  | Cynoglossus lighti | Cynoglossus lighti | Out |
|  | Paralichthydae | Paralichthys  olivaceus | Paralichthys  olivaceus | In |
|  |  | Pseudorhombus  pentophthalmus | Pseudorhombus  pentophthalmus | In |
|  |  | Tarphops  oligolepis | Tarphops  oligolepis | In |
|  | Pleuronectidae | Kareius bicoloratus | Platichthys  bicoloratus | In |
|  |  | Pleuronichthys  cornutus | Pleuronichthys  cornutus | In |
|  |  | Pleuronichthys  japonicus | Pleuronichthys  japonicus | In |
|  |  | Pseudopleuronectes  americanus | Pseudopleuronectes  americanus | Out |
|  |  | Pseudopleuronectes  yokohamae | Pseudopleuronectes  yokohamae | In |
|  | Soleidae | Aseraggodes  kobensis | Aseraggodes  kobensis | In |
|  |  | Heteromycteris  japonicus | Heteromycteris  japonicus | In |
|  |  | Pseudaesopia  japonica | Pseudaesopia  japonica | In |
| Tetraodontiformes | Monacanthidae | Rudarius ercodes | Rudarius ercodes | In |
|  |  | Stephanolepis  cirrhifer | Stephanolepis  cirrhifer | In |
|  |  | Thamnaconus  modestus | Thamnaconus  modestus | In |
|  | Tetraodontidae | Lagocephalus  spadiceus | Lagocephalus  spadiceus | In |
|  |  | Takifugu  alboplumbeus | Takifugu  alboplumbeus | In |
|  |  | Takifugu fasciatus x  Takifugu flavidus | NA | NA |
|  |  | Takifugu snyderi | Takifugu snyderi | In |
|  | Triacanthidae | Triacanthus  biaculeatus | Triacanthus  biaculeatus | In |
